# Accounting for technical noise in single-cell RNA sequencing analysis

**DOI:** 10.1101/116939

**Authors:** Cheng Jia, Derek Kelly, Junhyong Kim, Mingyao Li, Nancy R. Zhang

## Abstract

Recent technological breakthroughs have made it possible to measure RNA expression at the single-cell level, thus paving the way for exploring expression heterogeneity among individual cells. Current single-cell RNA sequencing (scRNA-seq) protocols are complex and introduce technical biases that vary across cells, which can bias downstream analysis without proper adjustment. To account for cell-to-cell technical differences, we propose a statistical framework, TASC (<u>T</u>oolkit for <u>A</u>nalysis of <u>S</u>ingle <u>C</u>ell RNA-seq), an empirical Bayes approach to reliably model the cell-specific dropout rates and amplification bias by use of external RNA spike-ins. TASC incorporates the technical parameters, which reflect cell-to-cell batch effects, into a hierarchical mixture model to estimate the biological variance of a gene and detect differentially expressed genes. More importantly, TASC is able to adjust for covariates to further eliminate confounding that may originate from cell size and cell cycle differences. In simulation and real scRNA-seq data, TASC achieves accurate Type I error control and displays competitive sensitivity and improved robustness to batch effects in differential expression analysis, compared to existing methods. TASC is programmed to be computationally efficient, taking advantage of multi-threaded parallelization. We believe that TASC will provide a robust platform for researchers to leverage the power of scRNA-seq.

## INTRODUCTION

Recent technological breakthroughs have made it possible to measure RNA expression at the single-cell level, thus paving the way for exploring gene expression heterogeneity among individual cells (1-4). The collection of abundances of all RNA species in a cell forms its “molecular fingerprint”, enabling the investigation of many fundamental biological questions beyond those possible by traditional bulk RNA sequencing experiments (5). With scRNA-seq data, one can better characterize the phenotypic state of a cell and more accurately describe its lineage and type.

Current scRNA-seq protocols are complex, often introducing technical biases that vary across cells (http://biorxiv.org/content/early/2015/08/25/025528), which, if not properly removed, can lead to severe type I error inflation in differential expression analysis. Compared to bulk RNA sequencing, in scRNA-seq the reverse transcription and preamplification steps lead to dropout events and amplification bias, the former describing the scenario in which a transcript expressed in the cell is lost during library preparation and is thus undetectable at any sequencing depth. In particular, due to the high prevalence of dropout events in scRNA-seq, it is crucial to account for them in data analysis, especially if conclusions involving low to moderately expressed genes are being drawn (7). In handling dropout events, existing studies take varying approaches: some ignore dropouts by focusing only on highly expressed genes (8,9), some model dropouts in a cell-specific manner (10-13), while others use a global zero-inflation parameter to account for dropouts (7).

Since each cell is processed individually within its own compartment during the key initial steps of library preparation, technical parameters that describe amplification bias and dropout rates should be cell-specific in order to adjust for the possible presence of systematic differences across cells. For example, a recent article by Leng *et al.* found significantly increased gene expression in cells captured from sites with small or large plate output IDs for data generated by the Fluidigm C1 platform (14). One way to quantify these biases, adopted by existing noise models (10-13), is to make use of spike-in molecules that comprise a set of external RNA sequences such as the commonly used external RNA Controls Consortium (ERCC) spike-ins (15), which are added to the cell lysis buffer at known concentrations (4,16). However, a challenge that cannot be ignored in the single-cell setting is that the wide range of concentrations of ERCC spike-ins makes it difficult to measure spike-ins with low concentrations, leading to the lack of reliable spike-in data for estimation of the dropout rates. For this reason, existing methods that model cell-specific dropout rates using spike-ins do not produce reliable estimates.

We propose here a new statistical framework that allows a more robust utilization of spike-ins to account for cell-specific technical noise. To obtain reliable estimates of cell-specific dropout parameters, we develop an empirical Bayes procedure that borrows information across cells. This is motivated by the observation that, although each cell has its own set of parameters for characterizing its technical noise, these parameters share a common distribution across cells which can be used to make the cell-specific estimates more stable. We demonstrate an application of this general framework by a likelihood-based test for differential expression. An advantage of the proposed framework over the existing approaches is that it can flexibly and efficiently adjust for cell-specific covariates, such as cell cycle stage or cell size, which may confound differential expression analysis.

## MATERIALS AND METHODS

### Data sets and pre-processing

<u>Zeisel</u> *<u>et al.</u>* <u>data:</u> scRNA-seq data from murine brain cells are acquired from Zeisel *et al.* (5). This data set, which employs UMIs, contains counts of 19,972 endogenous genes and 57 ERCC spike-ins of 3,005 cells from various regions of mouse brain. The cells are categorized into nine level-1 classes and 48 level-2 classes, with the level-2 classes considered relatively homogenous. In this paper, we focus our analyses on two level-2 classes, CA1Pyr1 and CA1Pyr2, which respectively contain 447 and 380 cells. The counts are preprocessed by selecting the top 25% of genes in total read account across the 827 cells, resulting in 6,405 genes in real data two-group comparison analysis. For studies involving class CA1Pyr2 only, selection of the top 25% of genes in the 447 cells yield 5,018 genes in the data set.

<u>SCAP-T data:</u> scRNA-seq data from murine brain cells are acquired from the SCAP-T study (dbGaP Study Accession phs000835.v4.p1). This data set, which does not have UMIs, contains counts of 46,422 endogenous genes and 87 ERCC spike-ins of 198 neurons and 26 astrocytes from mouse brain. The counts are preprocessed by two filtering procedures: Filter 1 keeps the top 25% of genes in total read account across all the cells. Filter 2 keeps all the genes with nonzero counts in 5 cells or more. Since neurons and astrocytes are processed on different days, this allows us to evaluate whether our model is able to capture and control batch effect.

### Modeling of technical variation

In scRNA-seq data, we have observed that the relationship between the mapped read count for a gene and its true expression level in a cell can be characterized using two functions, shown in Figure 1. For gene *g* in cell *c*, let *Z_cg_* be the indicator that dropout does *not* occur, i.e., that the gene is captured in the library. The probability of *Z_cg_* = 1 depends on the gene’s true absolute molecule count in the cell, denoted by *μ_cg._* We use a logistic model to capture this relationship

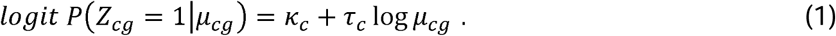

**Figure 1:**
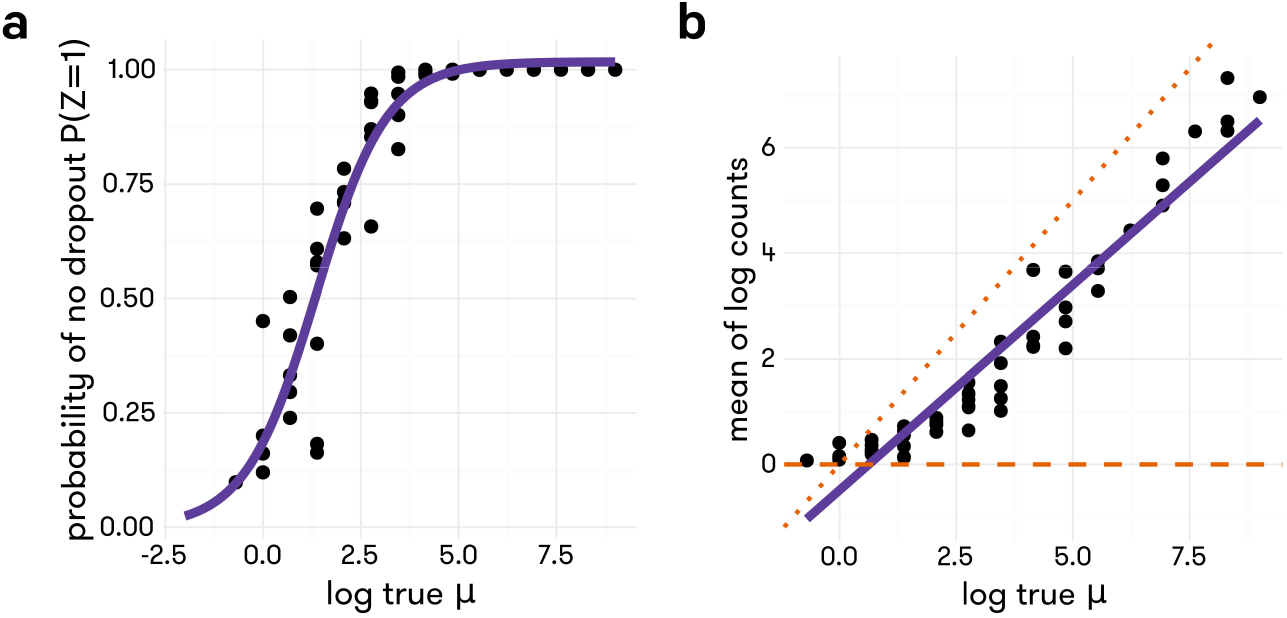
Proportion of cells with non-zero read count (in **a**) and mean across cells of log read count (in **b**) versus log true molecule count for ERCC spike-ins in Zeisel *et al.* data. Included in the plot are the best logistic curve fit (in **a**) and the best linear fit (in **b**).

For genes retained in the library (i.e. *Z_cg_ =* 1), the observed read count, denoted by *Y_cg_*, has expected value *λ_cg_* that increases linearly with *μ_cg_* on the log-log scale,

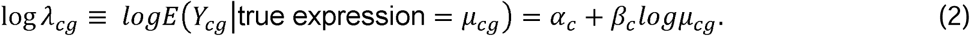

Figure 1 shows examples of the relationships depicted in (1) and (2) in the Zeisel *et al.* data (5); these relationships have also been seen in other studies (10,12). Note that the intercept *α_c_* is negative, indicating incomplete capture efficiency of reverse transcription, and that the slope, *β_c_*, when deviating from 1, reflects what is often called amplification bias. In experiments that use unique molecular identifiers (UMIs) (17), *Y_cg_* is the molecule count, and *βc* should be approximately 1. Together, functions (1) and (2) characterize the technical noise specific to each cell.

### Modeling of biological variation

The above observations have motivated the model shown in Figure 2, where the true but unobserved absolute expression level *μ_cg_* follows distribution *F_g_*, the specification of which depends on the analysis objective. For example, for the common task of detecting differentially expressed (DE) genes between groups, we assume *F_g_* follows a log-Normal distribution with mean *θ_gj_* and variance 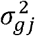, where *j* is the group identifier. The log-Normal distribution has been demonstrated previously to be a useful model for single cell gene expression (18), and lends computational simplicity to the estimation procedure. The technical noise in the cell is captured by the intermediate variables *Z_cg_*, characterized by (1), and *λ_cg_*, characterized by (2). Given *Z_cg_* and *λ_cg_*, the distribution of *Y_cg_* is 
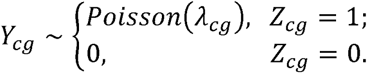

**Figure 2:**
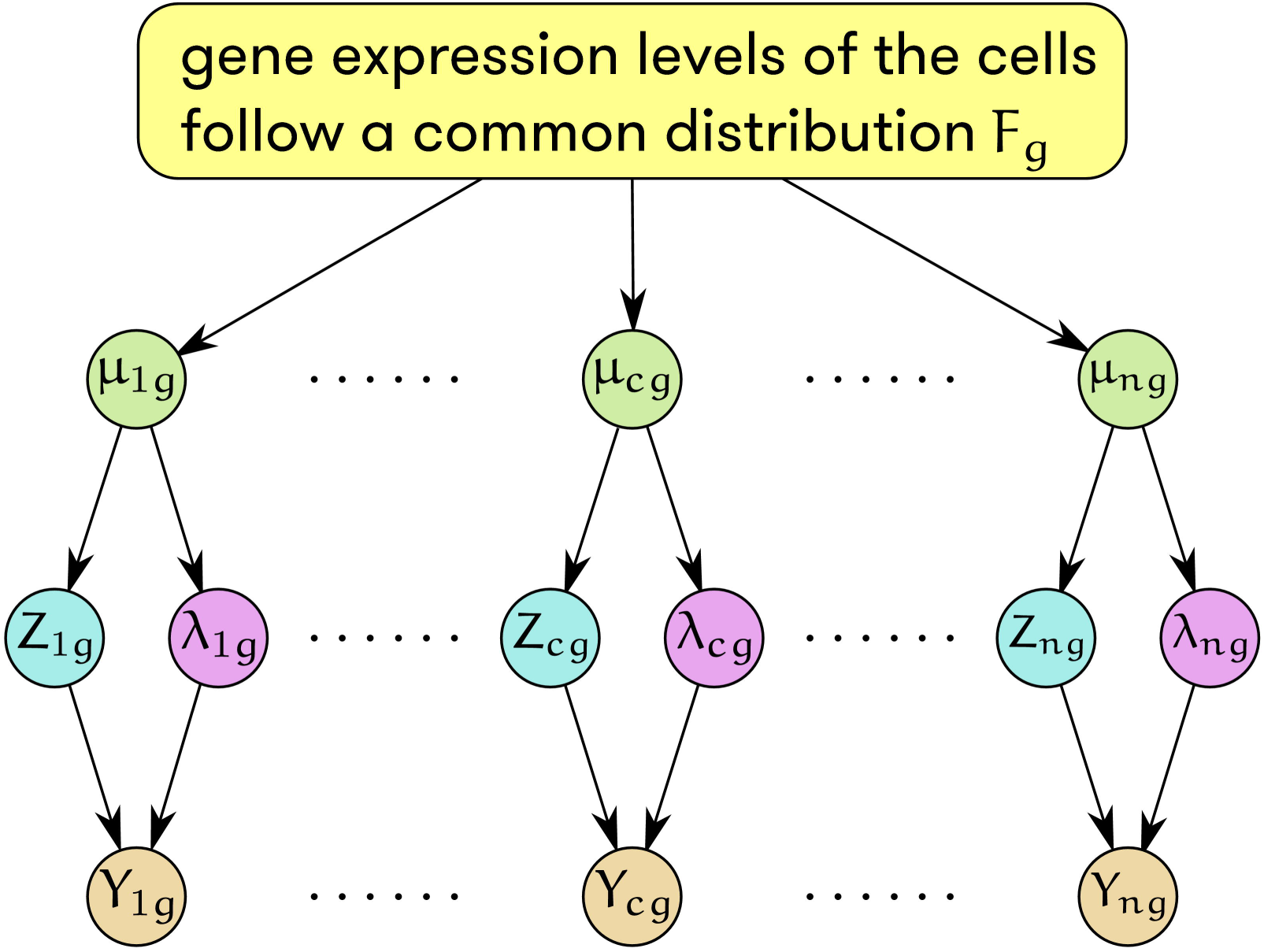
Schematic of TASC model for a single gene *g* across *n* cells, with *μ_cg_* being true absolute expression, *Y_cg_* being observed read count, and *Z_cg_, μ_cg_* being intermediate variables that model dropout and amplification, capture, and sequencing biases.

The Poisson distribution has been shown to be a reasonable approximation to the process of sampling reads from a fixed library, after removal of biological variation and technical biases (19).

### Estimation of cell-specific parameters describing technical variation

The cell-specific technical parameters (*α_c_*, *β_c_*, *κ _c_*, *τ_c_*) are estimated using ERCC spike-ins. Due to the lack of reliable spike-ins at low concentrations, we estimate (*κ_c_*, *τ_c_*) by empirical Bayes shrinkage. The cell-specific technical parameters, *α_c_* and *β_c_*, are estimated using simple linear regression with log*Y_cg_* as the response variable and the log of true amount of spiked-in molecules as the predictor variable, using only spike-ins that are detected in cell *c.* The dropout rate parameters, *κ_c_* and *τ_c_* are estimated using an empirical Bayes approach that can be summarized briefly into the following steps.

- Step 1: *κ_c_* and *τ_c_*, are estimated using logistic regression, logit 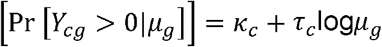, where *μ_g_* is the true expression level of spike-in *g* and it is assumed to be the same across all cells.
- Step 2: A bivariate normal distribution is fit to the estimated *κ_c_* and *τ_c_* across all cells to obtain the hyper-parameters, i.e., the mean and covariance matrix of *κ_c_* and *τ_c_*.
- Step 3: The estimated hyper-parameters are used to compute the posterior mean of *κ_c_* and *τ_c_*, 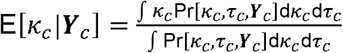 and 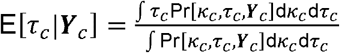, which will be used as the empirical Bayes estimates of *κ_c_* and *τ_c_*.

Refer to Supplementary Data for details on the computation of these estimates and the comparisons between empirical Bayes estimates and those derived from alternative approaches.

Figure 3 shows the distribution of estimated (*α_c_*, *β_c_*) and (*κ_c_*, *τ_c_*) across cells for the Zeisel data (5). The mean function, determined by (*α_c_*, *β_c_*), and the non-dropout rate function, determined by (*κ_c_*, *τ_c_*), are shown for four cells chosen to represent the middle and extremes of these distributions. The estimates of (*α_c_*, *β_c_*) have very small standard errors **(Figure S3)**, while empirical Bayes shrinkage greatly reduces the variance in the estimates of (*κ_c_*, *τ_c_*) (**Figure S4**). The presence of substantial variation in the magnitude of technical noise across cells underscores the need to account for such variation in downstream analyses.

**Figure 3:**
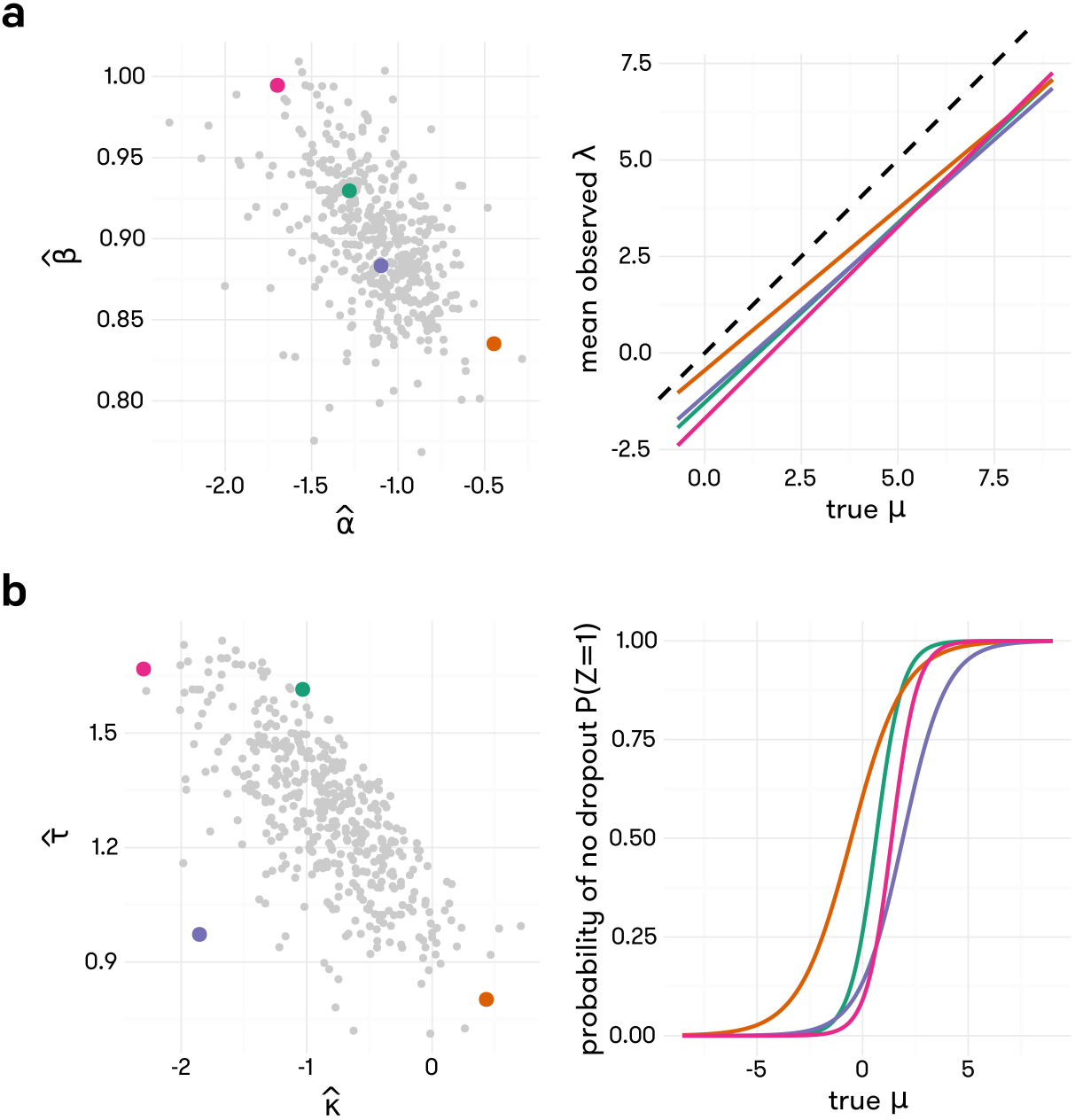
Distributions of empirically estimated values of (*α̂_c_*, *β̂_c_*) (*κ̂_c_*, *τ̂_c_*)and across all cells in Zeisel data. Four cells are selected from each plot to represent the distribution, and the line (in **a**) and logistic curve (in **b**) corresponding to the technical parameters estimated for these cells are shown in matching colors.

### Differential gene expression analysis

Previous studies have shown that cells vary in size, with larger cells having more RNA molecules to attain similar concentration levels to smaller cells (20). This indicates that to detect DE genes, it is more appropriate to test for concentration difference between groups. To allow this, we include cell size, which can be estimated by the ratio of reads from endogenous RNA to reads from spike-in sequences, as a covariate. Other potential covariates, such as cell cycle stage, can also be included in the model to avoid spurious association. For cell cycle, we add as covariate the expression of a curated set of marker genes, such as the set from Tirosh *et al.* (21), or a latent factor representing cell cycle, as in Buettner *et al.* (22). A likelihood-ratio test is developed to detect DE genes. Let *x_c_* be the group indicator for cell *c*, taking value 0 for group 1 and 1 for group 2, and let *U_c_* be the optional vector of covariates corresponding to cell *c.* The true expression of a given gene *g* in cell *c* is assumed to have a log-Normal distribution with mean 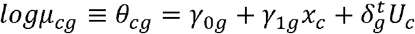, where *γ*_1*g*_, the between-group difference in log mean, is the parameter of interest. To determine whether gene *g* is differentially expressed, we test *H*_0_:*γ*_1*g*_ = 0 VS *H*_1_:*γ*_1*g*_ ≠ 0. Let 𝓛̂_0*g*_ and 𝓛̂_1*g*_ be the maximized likelihoods achieved under the null and alternative hypotheses, respectively. The likelihood ratio test statistic for gene *g* is *T̂_g_* = 2[𝓛̂_1*g*_ − 𝓛̂_0*g*_]. Under the null hypothesis, this test statistic approximately follows a chi-squared distribution with one degree of freedom. The parameters in this model, including the variance *σ_g_*^2^ of the log-Normal distribution, can be estimated by numerical optimization or by a scalable Expectation-Maximization (EM) algorithm if the number of covariates is large. Please refer to the **Supplementary Data** for details.

## RESULTS

In this section, we evaluate the performance of TASC on both simulated and two real scRNA-seq data sets and compare it with four existing methods, including SCDE (10), MAST (11), and DESeq2 (23), and SCRAN (24). As SCRAN only provides normalized read counts, we perform differential expression analysis using DESeq2 with SCRAN normalized read counts. We include two versions of SCRAN in our evaluation, the original SCRAN, and SCRAN.SP that utilizes ERCC spike-ins in normalization. These methods are rated in terms of type I error rate and power in detecting DE genes, and their results on a real data set with genuine gene expression difference.

### Type I error rates in the absence of batch effects

To assess the accuracy of type I error control of TASC and other existing methods, 447 cells from the level-2 class “CA1Pyr2” from the Zeisel *et al.* data, which is the largest level-2 class, are randomly split into two groups of roughly equal size. Differential expression analyses are performed with TASC, SCDE, MAST, DESeq2, SCRAN and SCRAN.SP. Raw p-values are extracted from each method, and the performance of each method is assessed by histograms and quantile-quantile plots of the corresponding p-values, shown in Figure 4. Our results show that TASC, DESeq2, SCRAN and SCRAN.SP have p-values that are uniformly distributed as expected under the null, whereas SCDE is overly conservative with enrichment of p-values near one, and MAST is severely anti-conservative with enrichment of p-values near zero.

**Figure 4:**
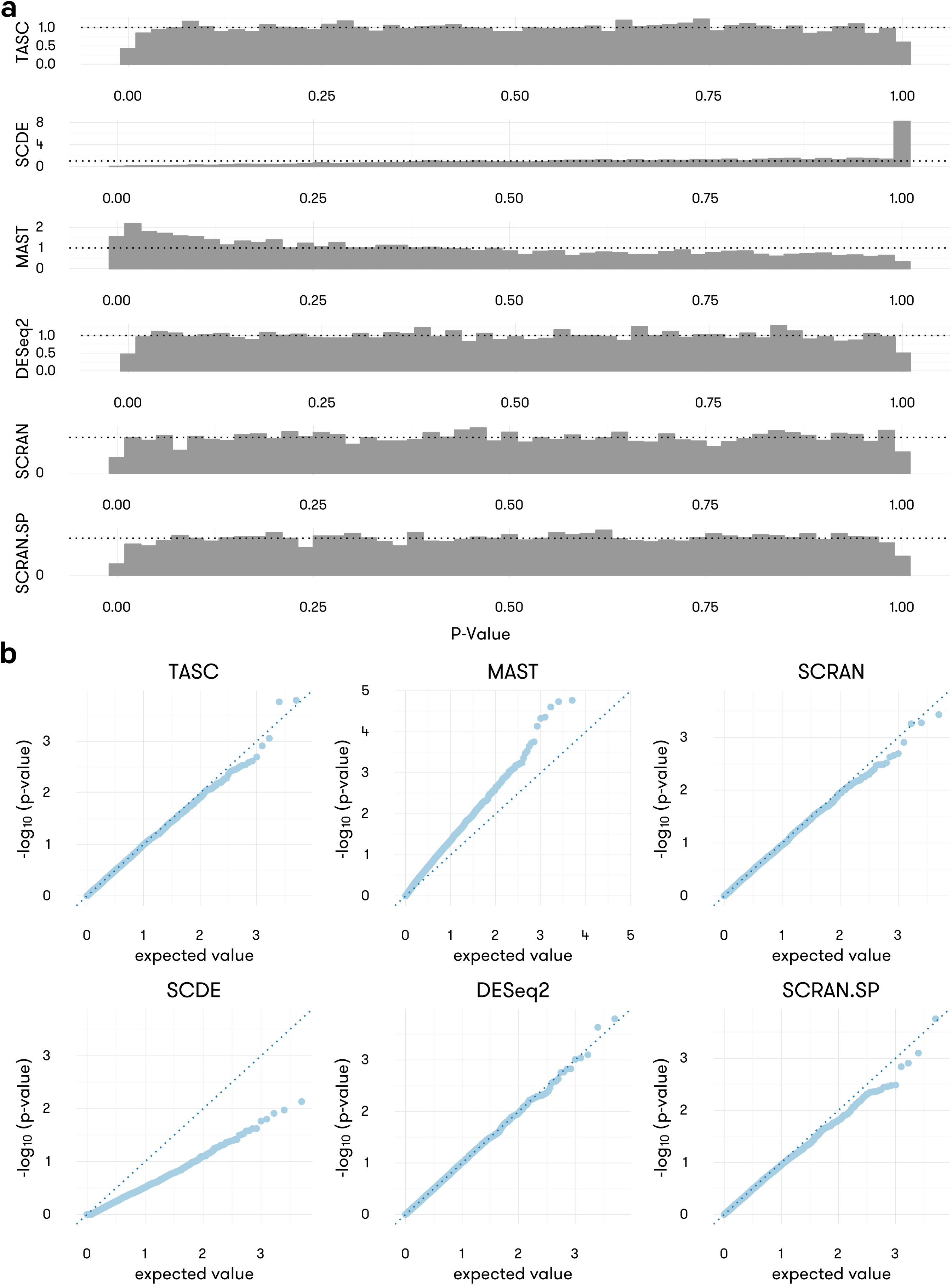
Distribution of achieved p-values (in **a**) and the corresponding quantile-quantile plots (in **b**) for four methods applied to CA1Pyr2 cells from Zeisel *et al.* data, split randomly into two groups, thus emulating a case where all p-values should be uniformly sampled from [0,1].

### Type I error rates in the presence of batch effects

Batch effects are common in scRNA-seq data

(http://biorxiv.org/content/early/2015/08/25/025528). To evaluate effectiveness in type I error control in the presence of batch effects, we have generated a data set that contains batch effects as characterized by systematic differences in the technical parameters (*α_c_*, *β_c_*, *κ_c_*, *τ_c_*) between groups. To introduce batch differences between the two groups under comparison, cell-specific technical parameters (*α_c_*, *β_c_*) and (*κ_c_*, *τ_c_*) are estimated from the cells in “CA1Pyr2” class and a bivariate normal distribution is fit separately to {(*α_c_*, *β_c_*)} and to {(*κ_c_*, *τ _c_*)}. One group in the simulated data draws its cell-specific technical parameters from these empirical distributions, and the other group draws its technical parameters from distributions where the mean(s) of combinations of technical parameters are shifted by amounts shown on the axes of the heatmaps in Figure 5. The magnitude of the shift represents the severity of batch effect difference between the two groups. The rest of the parameters controlling the expression of genes are the same for the two groups and are derived from estimates from the “CA1Pyr2” class. Simulations are performed to generate the counts of 5,018 genes in 100 cells (50 in each group). Differential expression analyses are performed and the raw p-values are used to estimate the false positive rate (FPR). The deviation of the estimated FPR from the expected value is plotted on heatmaps to reflect the type I error rates under varying severity of batch effects. Figure 5 shows that TASC has well controlled type I error rates across a wide range of batch effect severity, whereas SCDE appears to be conservative overall, and MAST, DESeq2, SCRAN and SCRAN.SP are anti-conservative and susceptible to batch effects.

**Figure 5:**
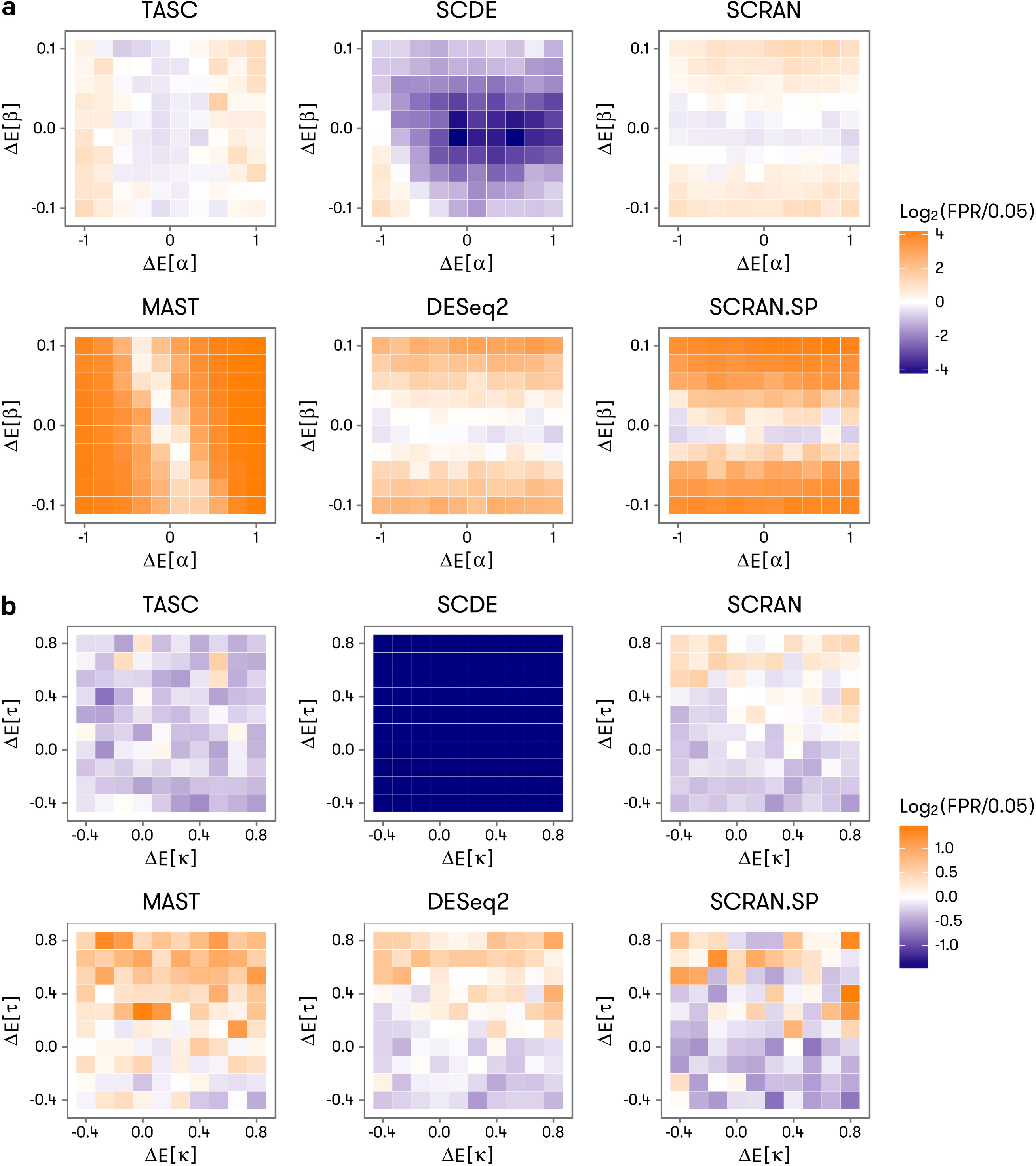
Accuracy of false positive rate control under mild to severe batch effects for TASC, SCDE, MAST, and DESeq2. The batch effect severity takes the form of between-group difference in the expected values of the technical parameters, denoted by Δ *E*[*τ*] and Δ *E*[*κ*] (in **a**), and Δ *E*[*α*] and Δ *E*[*β*] (in **b**) in the axes of the heatmaps. The color scale of the heatmaps reflects deviation of achieved false positive rate from the target value of 0.05 used in the tests.

To compare the methods with regards to their type I error rate under a real data scenario, we analyzed the SCAP-T data, which includes astrocytes and neurons that were processed on different days. This data set provides a perfect example to illustrate the impact of batch effect. As shown in **Supplementary Figure S23** the technical parameters, as characterized by (*α_c_*, *β_c_*, *κ_c_*, *τ_c_*), can clearly capture the batch effect. To assess whether type I error is controlled under the null scenario, it is necessary to compare two groups of cells that are of the same type. To perform this assessment, we divide the 198 neurons into two groups in which group 1 includes the 26 neurons that have similar (*α_c_*, *β_c_*, *κ_c_*, *τ _c_*) values as the astrocytes and group 2 includes the remaining 172 neurons. The methods TASC, SCDE, MAST, DESeq2, SCRAN, and SCRAN.SP are then applied to these two groups, and the proportion of genes reported to be DE is reported in Table 1. We see that TASC has well controlled type I error rates at all assessed significance levels, whereas all other methods (SCDE, MAST, DESeq2, SCRAN, and SCRAN.SP) have severely inflated type I error rates, especially when the p-value threshold is reduced to 0.001 and 0.0001. For example, consider DESeq2, which, according to our simulations, has well-controlled type I error when there are no batch effects. At significance level of 0.001, DESeq2 has false positive rate of 1.7%, a 17-fold inflation, and at significance level of 0.0001, DESeq2 has false positive rate of 0.76%, corresponding to a 76-fold inflation. Even SCDE, which tends to be conservative when there are no batch effects, suffer from type I inflation in this real data scenario that contains a possible batch effect. The patterns are similar when we consider all genes in the evaluations.

**Table 1:** Proportion of DE genes identified by each method in SCAP-T data at varying significance levels. Filter 1 keeps the top 25% of genes in total read account across all the cells. Filter 2 keeps all the genes with non-zero counts in 5 cells or more. Naive SCRAN without the use of spike-ins is not included in this comparison, for the package fails to run due to there being “not enough cells in each cluster for specified ‘sizes’”.

| *Filter* | **Filter 1** |  |  |  |  | **Filter 2** |  |  |  |  |
| --- | --- | --- | --- | --- | --- | --- | --- | --- | --- | --- |
| *Significance Level* | **0.05** | **1e-2** | **1e-3** | **1e-4** | **1e-5** | **0.05** | **1e-2** | **1e-3** | **1e-4** | **1e-5** |
| *TASC* | 2.15e-2 | 2.33e-3 | 0 | 0 | 0 | 3.09e-2 | 5.78e-3 | 5.36e-4 | 8.93e-5 | 2.98e-5 |
| *SCDE* | 4.28e-2 | 1.65e-2 | 4.74e-3 | 1.46e-3 | 3.45e-4 | 9.42e-2 | 3.46e-2 | 8.72e-3 | 1.79e-3 | 1.79e-4 |
| *MAST* | 4.89e-2 | 1.12e-2 | 1.81e-3 | 7.75e-4 | 2.58e-4 | 7.42e-2 | 2.62e-2 | 1.21e-2 | 8.45e-3 | 6.25e-3 |
| *DESeq2* | 1.70e-1 | 8.44e-2 | 3.52e-2 | 1.69e-2 | 8.27e-3 | 1.25e-1 | 5.91e-2 | 2.32e-2 | 1.01e-2 | 4.79e-3 |
| *SCRAN.SP* | 2.50e-1 | 1.39e-1 | 6.63e-2 | 3.67e-2 | 2.11e-2 | 1.69e-1 | 9.13e-2 | 4.19e-2 | 2.21e-2 | 1.21e-2 |

### Evaluation of power

To investigate the power of the methods under realistic scenarios, we continue to utilize the 5,018 genes from the “CA1Pyr2” class in Zeisel *et al.* data set. Among them, 4,018 genes are designated as “true non-DE”, whose counts are directly extracted from the Zeisel *et al.* data set after group membership randomization. The remaining 1,000 are designated as “true DE”, whose counts are simulated from parameters estimated with real data, with an induced between-group fold change that is randomly sampled from a distribution that generates more genes with weak to moderate expression difference than strong difference (details are given in Section 2.3 of **Supplementary Data**). DE analyses are performed with all methods, and raw p-values are used to estimate the power of each method at various significance levels. The average power curves in Figure 6A are obtained by smoothing the estimated power across genes with similar fold change. Our results demonstrate that TASC has the highest power, followed by SCRAN.SP, SCRAN, DESeq2, MAST, and SCDE. Figure 6B shows that the higher sensitivity of TASC is more pronounced when fold change is moderate; for example, when fold change is 1.75, at the 0.0001 significance level, the average power of TASC is 8%, 20%, 25%, 37%, and 428% higher than SCRAN.SP, SCRAN, DESeq2, MAST, and SCDE, respectively.

**Figure 6:**
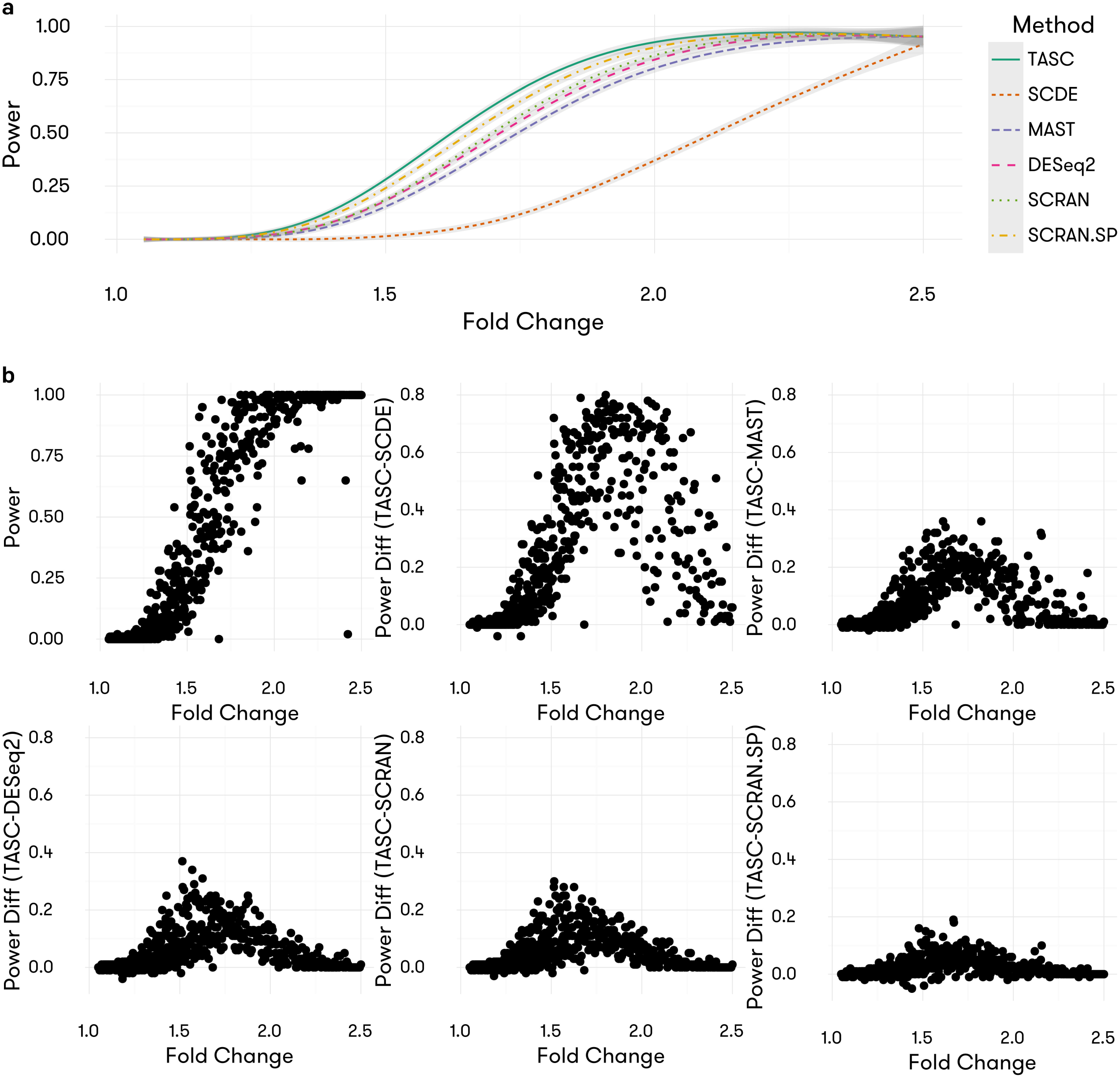
Achieved power of TASC, SCDE, MAST, DESeq2, SCRAN, and SCRAN.SP for detecting varying fold changes in mean in the simulated data set within 100 cells in each group. Results both with (SCRAN.SP) and without (SCRAN) the use of ERCC are included for SCRAN. **b**) Power differences between TASC and the other methods in the simulated data set. Details of data simulation, and power comparisons under more comprehensive settings, are in Supplementary Data.

The analysis also indicates the importance of sample size in DE analysis in scRNA-seq (**Supplementary Figures S21-22**).

### Differential Expression analysis on real data

We continue to use the Zeisel *et al.* data set to evaluate methods performance when true differential expression is present. All five methods are applied to detect DE genes between the two level-2 classes “CA1Pyr2” (*n* = 380 randomly chosen from 447 cells) and “CA1Pyr1” (*n* = 380). Since these two level-2 classes represent different cell type groups, we expect genuine gene expression differences between them. To evaluate the impact of sample size, the two groups are subsampled to 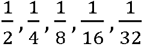 of their original size, and differential expression analyses are performed on each subsampled data set. The raw p-values are used to detect DE genes at the 0.0001 significance level, and the number of detected DE genes is plotted against the sample size for each method. The numbers of detected DE genes are shown in Figure 7. Consistent with our simulations, SCDE finds the least number of DE genes, followed by MAST, whereas SCRAN.SP detects the most number of DE genes when *n* is greater than 100. TASC, SCRAN, and DESeq2 detect similar number of DE genes across most sample sizes.

**Figure 7:**
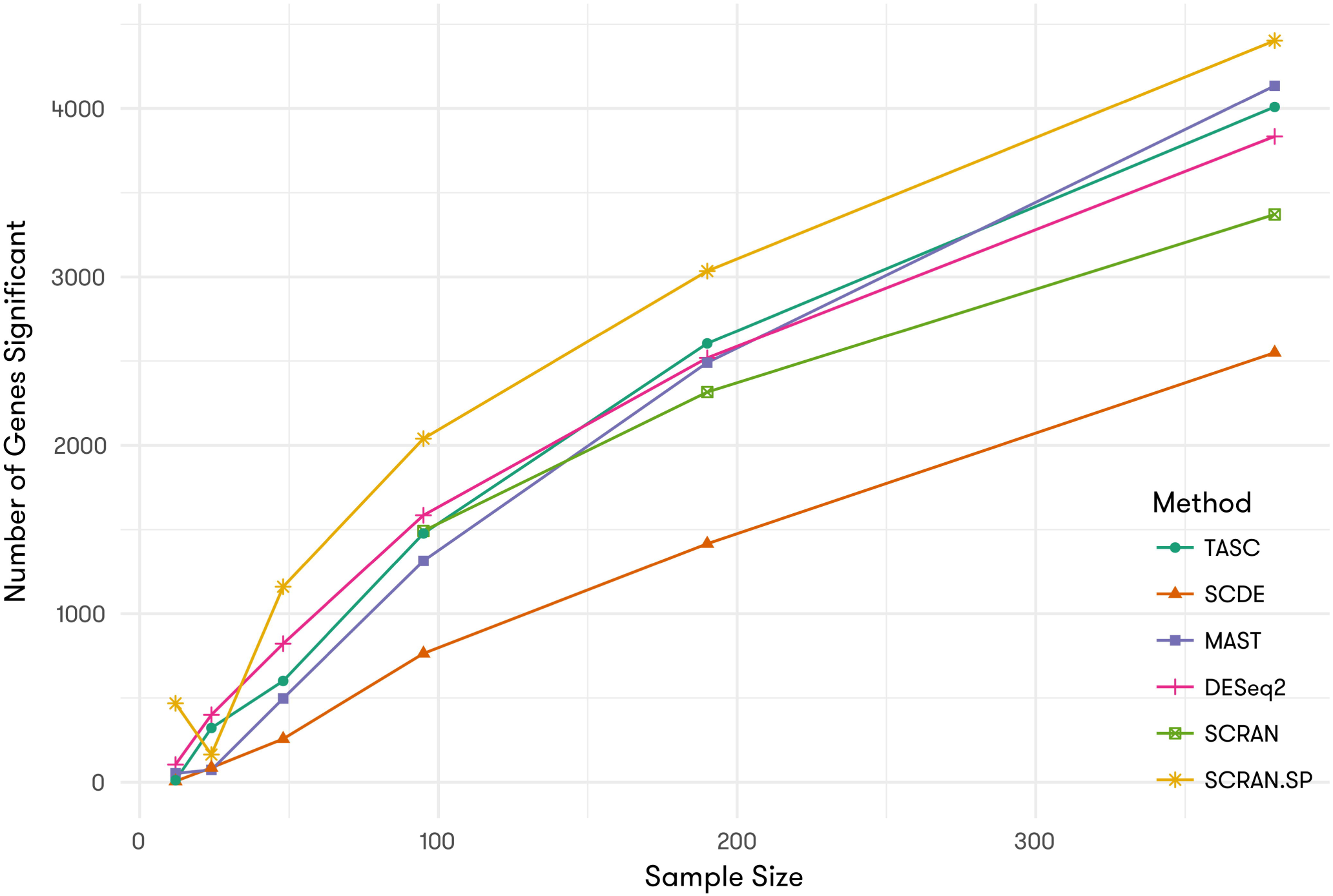
Number of DE genes identified by each method between two level-2 classes in Zeisel *et al.* data at the 0.0001 significance level, under varying sample sizes.

We further evaluate the overlap of DE genes detected by different methods. Since we do not know what genes are truly differentially expressed, for each method, we calculate the percentage of DE genes that are unique to that method. This is based on the rationale that genes detected by two or more methods are more likely to be true DE. Subsequently, a better performing method shall have a larger number of detected genes but a smaller percentage of genes uniquely detectable by itself. In the full data set, only 0.8% and 1.12% of the DE genes detected by SCRAN and TASC are unique to themselves. In contrast, the corresponding percentage increases to 1.9% for DESeq2, 20.94% for SCRAN.SP, 1.74% for MAST, and 2.3% for SCDE. The higher percentages for these three methods indicate possible false positive results. Similar patterns have been observed for comparisons with smaller sample size.

## DISCUSSION

The advent of scRNA-seq has made it possible to explore cellular heterogeneity with unprecedented resolution. For the first time, we are able to measure the cell-to-cell variation of RNA expression of all genes in the genome across hundreds to thousands of cells. However, due to the limitations of current technology, scRNA-seq data are often noisy. Failure to account for technical noise can lead to biased downstream analyses and misleading results. To take full advantage of scRNA-seq, it is crucial to account for technical noise so as to better quantify biological variation. Here we have described a statistical framework, TASC, that accurately estimates cell-specific technical biases, adjusts for them in differential expression analysis, and consequently produces results that are more robust to batch effects that exhibit as systematic differences between cells.

TASC utilizes information in spike-ins to account for technical noise in a cell-specific manner. Compared to the traditional bulk RNA sequencing, in scRNA-seq the reverse transcription and preamplification steps can lead to pervasive dropout events and amplification bias. While amplification bias can be alleviated by the use of UMIs, dropout events are harder to control. To reliably estimate cell-specific dropout parameters under the paucity of reliable spike-ins at low concentrations, we have developed an empirical Bayes procedure that borrows information across cells. The accuracy of this empirical Bayes procedure has been examined in simulations based on real scRNA-seq data.

Our evaluations show that TASC is always slightly conservative. However, we are willing to accept this slight conservativeness, since the data used in our evaluations are generated under an ideal null distribution, and we believe it is more meaningful to examine each method’s performance in noisier data where strong deviations from the null can be observed. This is the motivation for our analysis of the data set from SCAP-T involving the comparisons of two groups of neurons with batch effects. Our results show that TASC achieves accurate type I error control under this noisy setting, whereas other methods have substantially inflated type I errors.

Since the performance of TASC relies on spike-ins, it is important to determine if the spike-in data are reliable so that the algorithm can give reasonable results. Before applying TASC, we would always recommend a visual examination of the spike-in data using plots similar to Figure 1. One should see the sigmoidal curve for the probability of dropout, and a line for the log mean observed versus log mean actual counts. These curves, which are computed across cells, give a sanity check that the spike-ins are diluted to an appropriate concentration. For each cell, one could also plot the log actual count versus the log observed count of the spike-in molecules, and compute the *R*^2^ of the linear fit (**Supplementary Figures S24** and **S25**). If this *R*^2^ is too low, the spike-in data for that cell may give poor estimates of technical parameters. One could throw away cells with *R*^2^ values below a threshold, on the basis of unreliable spike-ins, but we believe that is not necessary due to the shrinkage imposed by our empirical Bayes model. For those cells where the observed spike-in counts are too noisy, estimates of the technical parameters are shrunk more aggressively to the cross-cell means, and thus for these less-than-ideal cases one could still have usable technical parameters for normalization.

In scRNA-seq data analysis, it is critical to filter out cells with low quality. A commonly used strategy for filtering is to remove cells in which more than 20% reads come from ERCC spike-ins, on the basis that these cells may have been compromised during the experiment. We are not against this filtering strategy, however, we believe that this pre-filtering is not strictly necessary if the adjustment is made for technical dropout and cell size in downstream analyses. Since the cells with >20% ERCC reads would have small estimated size, and since most genes in a small cell would have zero count or small count prone to dropout, the zeros in such a cell would no longer be outliers in a differential expression analysis. Hence, by careful modeling of technical bias and by allowing for cell size as a covariate, TASC reduces the contribution of these low quality cells to the analysis and avoids the use of an arbitrary cutoff in eliminating cells.

An important feature of TASC is the ability to adjust for covariates such as cell size and cell cycle. **Supplementary Figures S26** and **S27** show the cell size histogram for the two data sets we analyzed. For the Zeisel *et al.* data, since the two groups being compared are both pyramidal neurons, cell sizes are comparable. But for the SCAP-T data, astrocytes and neurons differ substantially in cell size. If the goal is to compare astrocytes and neurons, then adjustment of cell size might be necessary. In practice, when should cell size adjustment be made? This depends on the biological question: If the goal is to find genes that differ in concentration between two cell types, then one should adjust for cell size. If one doesn’t adjust for cell size, then most genes would be significant, since the expression of most genes scale with cell size, and thus, the genes that are markers for real pathway differences between cell-types would be hard to detect. Ultimately, whether to adjust for cell size is a decision for the user, and our goal through TASC is to provide the flexibility. For cell cycle, we rely on a curated set of marker genes, such as the set from Tirosh *et al.* (21), or a latent factor representing cell cycle, as from Buettner *et al.* (22). The two data sets that we analyze involve non-cycling cells, and thus adjustment for cell cycle is not necessary.

The hierarchical mixture model underlying TASC allows for flexible modeling of the true biological variation of gene expression across cells, and thus can be adapted to tackle many interesting biological questions. For example, ranking the estimated values of *σ_g_^2^* allows us to identify biologically variable genes. The posterior expectation of *μ_cg_* also gives us the inferred true expression value given the observed read counts. To illustrate the importance of accurate adjustment for cell-specific technical noise, in this paper we benchmark TASC against existing methods for differential expression analysis. TASC currently makes the assumption that true expression levels follow log-Normal distributions, however it can be readily extended to assume other forms for *F_g_*, for example, the Poisson-Beta distribution if the modeling of transcriptional bursting is of interest (25).

TASC incorporates the estimated technical parameters, which reflect cell-to-cell differences that may lead to batch effects, into a hierarchical mixture model to estimate the biological variance of a gene and to detect DE genes. The EM algorithm implemented in TASC offers a flexible and efficient approach to adjust for additional covariates to further eliminate confounding originated from cell size and cell cycle differences. In our evaluations, TASC appears to be robust in the detection of DE genes when batch effects are present.

The current implementation of TASC assumes that the true expression levels of a gene follow a logNormal distribution in cells. We recognize that logNormal does not entirely reflect the true distribution as transcriptional bursting could lead to zeros in the expression. A more realistic distribution is zero-inflated logNormal, which accounts for true zeros in gene expression. We choose to use logNormal because it is simple and fast to fit. To examine the impact of misspecification of distribution, we have conducted additional simulations and found that the use of the simplified logNormal distribution does not lead to noticeable drop in power when the goal is to detect mean expression difference between two groups. However, a zero-inflated distribution for true expression distribution allows the detection of subtler signals beyond change in mean, an analysis goal that was emphasized in Vallejos *et al.* (26). Examples of such subtle but important signals include change in the probability of a gene having nonzero expression, and change in variance of the nonzero mixture component. We are currently extending TASC to account for this possibility.

TASC is implemented in an open-source program (https://github.com/scrna-seq/TASC), with multithreading acceleration by openMP. For example, a data set of 104 cells and 6,405 genes takes 45MB of memory and 18.6 minutes using 20 cores (Intel(R) Xeon(R) CPU E5-2660 v3 @ 2.60GHz) with Laplacian approximation using the binary we provided. Better performance can be achieved when using binaries compiled on the user’s hardware. We believe that TASC will provide a robust platform for researchers to leverage the power of scRNA-seq.

## SUPPLEMENATARY DATA

Supplementary Data are available at NAR Online.

## ACKNOWLEDGEMENT

The authors thank Drs. James Eberwine and Arjun Raj for helpful discussions.

## ACCESSION NUMBER

The SCAP-T sequencing data have been deposited in dbGap (accession number phs000835.v4.p1).

## References

1. Sandberg, R. (2014) Entering the era of single-cell transcriptomics in biology and medicine. Nat Methods, 11, 22–24.

2. Kolodziejczyk, A.A., Kim, J.K., Svensson, V., Marioni, J.C. and Teichmann, S.A. (2015) The technology and biology of single-cell RNA sequencing. Mol Cell, 58, 610–620.

3. Eberwine, J., Sul, J.Y., Bartfai, T. and Kim, J. (2014) The promise of single-cell sequencing. Nature methods, 11, 25–27.

4. Bacher, R. and Kendziorski, C. (2016) Design and computational analysis of single-cell RNA-sequencing experiments. Genome Biol, 17, 63.

5. Zeisel, A., Munoz-Manchado, A.B., Codeluppi, S., Lonnerberg, P., La Manno, G., Jureus, A., Marques, S., Munguba, H., He, L., Betsholtz, C. et al. (2015) Brain structure. Cell types in the mouse cortex and hippocampus revealed by single-cell RNA-seq. Science, 347, 1138–1142.

6. Leng, N., Chu, L.F., Barry, C., Li, Y., Choi, J., Li, X., Jiang, P., Stewart, R.M., Thomson, J.A. and Kendziorski, C. (2015) Oscope identifies oscillatory genes in unsynchronized single-cell RNA-seq experiments. Nat Methods, 12, 947–950.

7. Pierson, E. and Yau, C. (2015) ZIFA: Dimensionality reduction for zero-inflated single-cell gene expression analysis. Genome Biol, 16, 241.

8. Shalek, A.K., Satija, R., Adiconis, X., Gertner, R.S., Gaublomme, J.T., Raychowdhury, R., Schwartz, S., Yosef, N., Malboeuf, C., Lu, D. et al. (2013) Single-cell transcriptomics reveals bimodality in expression and splicing in immune cells. Nature, 498, 236–240.

9. Shalek, A.K., Satija, R., Shuga, J., Trombetta, J.J., Gennert, D., Lu, D., Chen, P., Gertner, R.S., Gaublomme, J.T., Yosef, N. et al. (2014) Single-cell RNA-seq reveals dynamic paracrine control of cellular variation. Nature, 510, 363–369.

10. Kharchenko, P.V., Silberstein, L. and Scadden, D.T. (2014) Bayesian approach to single-cell differential expression analysis. Nature methods, 11, 740–742.

11. Finak, G., McDavid, A., Yajima, M., Deng, J., Gersuk, V., Shalek, A.K., Slichter, C.K., Miller, H.W., McElrath, M.J., Prlic, M. et al. (2015) MAST: a flexible statistical framework for assessing transcriptional changes and characterizing heterogeneity in single-cell RNA sequencing data. Genome Biol, 16, 278.

12. Kim, J.K., Kolodziejczyk, A.A., lllicic, T., Teichmann, S.A. and Marioni, J.C. (2015) Characterizing noise structure in single-cell RNA-seq distinguishes genuine from technical stochastic allelic expression. Nature communications, 6, 8687.

13. Vallejos, C.A., Marioni, J.C. and Richardson, S. (2015) BASiCS: Bayesian Analysis of Single-Cell Sequencing Data. PLoS Comput Biol, 11, el004333.

14. Leng, N., Choi, J., Chu, L.F., Thomson, J.A., Kendziorski, C. and Stewart, R. (2016) OEFinder: a user interface to identify and visualize ordering effects in single-cell RNA-seq data. Bioinformatics, 32, 1408–1410.

15. Jiang, L., Schlesinger, F., Davis, C.A., Zhang, Y., Li, R., Salit, M., Gingeras, T.R. and Oliver, B. (2011) Synthetic spike-in standards for RNA-seq experiments. Genome research, 21, 1543–1551.

16. Stegle, O., Teichmann, S.A. and Marioni, J.C. (2015) Computational and analytical challenges in single-cell transcriptomics. Nat Rev Genet, 16, 133–145.

17. Islam, S., Zeisel, A., Joost, S., La Manno, G., Zajac, P., Kasper, M., Lonnerberg, P. and Linnarsson, S. (2014) Quantitative single-cell RNA-seq with unique molecular identifiers. Nature methods, 11, 163–166.

18. Bengtsson, M., Stahlberg, A., Rorsman, P. and Kubista, M. (2005) Gene expression profiling in single cells from the pancreatic islets of Langerhans reveals lognormal distribution of mRNA levels. Genome research, 15, 1388–1392.

19. Grun, D., Kester, L. and van Oudenaarden, A. (2014) Validation of noise models for single-cell transcriptomics. Nature methods, 11, 637–640.

20. Padovan-Merhar, O., Nair, G.P., Biaesch, A.G., Mayer, A., Scarfone, S., Foley, S.W., Wu, A.R., Churchman, L.S., Singh, A. and Raj, A. (2015) Single mammalian cells compensate for differences in cellular volume and DNA copy number through independent global transcriptional mechanisms. Mol Cell, 58, 339–352.

21. Tirosh, I., Izar, B., Prakadan, S.M., Wadsworth, M.H., 2nd, Treacy, D., Trombetta, J.J., Rotem, A., Rodman, C., Lian, C., Murphy, G. et al. (2016) Dissecting the multicellular ecosystem of metastatic melanoma by single-cell RNA-seq. Science, 352, 189–196.

22. Buettner, F., Natarajan, K.N., Casale, F.P., Proserpio, V., Scialdone, A., Theis, F.J., Teichmann, S.A., Marioni, J.C. and Stegle, O. (2015) Computational analysis of cell-to-cell heterogeneity in single-cell RNA-sequencing data reveals hidden subpopulations of cells. Nat Biotechnol, 33, 155-160.

23. Love, M.I., Huber, W. and Anders, S. (2014) Moderated estimation of fold change and dispersion for RNA-seq data with DESeq2. Genome Biol, 15, 550.

24. Lun, A.T., Bach, K. and Marioni, J.C. (2016) Pooling across cells to normalize single-cell RNA sequencing data with many zero counts. Genome Biol, 17, 75.

25. Raj, A., Peskin, C.S., Tranchina, D., Vargas, D.Y. and Tyagi, S. (2006) Stochastic mRNA synthesis in mammalian cells. PLoS Biol, 4, e309.

26. Vallejos, C.A., Richardson, S. and Marioni, J.C. (2016) Beyond comparisons of means: understanding changes in gene expression at the single-cell level. Genome Biol, 17, 70.

